# Identification of the potential function-specific sites in subunits of vertebrate neuronal nicotinic acetylcholine receptors

**DOI:** 10.1101/2024.05.09.593419

**Authors:** Zhouhai Zhu, Fengyu Zhang, Ying Guan, Zhenhua Pan, Meng Li, Ju Wang

## Abstract

The nicotinic acetylcholine receptors (nAChR) are complexes that are composed of subunits evolved from common ancestor. Although the subunits are similar in sequences and structures, their molecular function varies significantly. Therefore, detecting the molecular sites specific to each subunit is important to understand the property of the subunits and the receptors formed by them. As we know, the molecular sites critical to the structure and molecular function of a protein family usually are conserved in evolution, and those specific to each member of the family are often closely related to its structural and functional specificity. In this study, we analyzed the sequence specific sites in nAChR subunits by adopting the evolutionary trace method and the two-state model, and explored the relationship between structure and function in combination with the spatial location of the sites. The results showed that the detected sites in α7 subunit were closely related to ligand binding and conformational changes of the energetic coupling pathway. The conserved sites tended to be distributed in the interior of the spatial structure of protein molecules, and the sites potentially related to new functions were distributed on the surface of the spatial structure. In summary, our results could be helpful to understanding the molecular features related to the function specificity and diversity of the nAChR subunits.

## Introduction

Acetylcholine (ACh) is a neurotransmitter that plays a wide variety of roles in the brain and other organs [1–2]. In the nervous system, ACh can modulate numerous physiological functions by binding to the nicotinic acetylcholine receptors (nAChRs). Structurally, nAChRs belong to the super-family of Cys-loop ligand-gated ion channels that comprise five subunits surrounding a central channel [3]. The nAChRs are diverse in properties, which can be attributed to the large number of possible combinations of the subunits. In the vertebrate genomes, seventeen nAChR subunits have been identified so far, with the α2-α7 and β2-β4 subunits commonly known as the neuronal nAChR subunits [4]. These subunits share a common architecture, featuring an extracellular amino-terminal domain housing the ACh binding sites that faces the synaptic gap, followed by three hydrophobic transmembrane segments (M1–M3), a sizable intracellular loop, and finally a fourth hydrophobic transmembrane region (M4)[5]. In each subunit, the amino acid residues involved in ACh binding can be clustered into short segments called loops A, B, and C (the primary component) and D, E, and F (the complementary component). Among these loops, A, B, D, and F, are positioned within the center of the interface of two neighboring subunits, forming a hydrophobic cavity for nicotinic ligands to bind [6]. Upon binding to nAChRs, the agonist induces a conformation change in the ion channel, leading to the opening of the central pore and facilitating the transient passage of ions for a brief duration. Subsequently, the channel transitions to a non-conductive state [2].The extracellular domain (ECD) and the transmembrane channel are connected by the so-called energetic coupling pathway (ECP) that comprises the Cys-loop, β8-β9 loop and TM2-TM3 loop, which enables the transmission of conformational alterations from the ECD to the ion gating portion.

Although the sequence and structure of nAChR subunits are similar, they usually have different properties. For example, ligand-binding sites can only be found on subunits like α2-α4, α6 or α7, while the functional and pharmacological properties of the receptors are dependent on subunits like α5 and β3 [7–9]. The nAChR subunits can form either homo- or hetero- pentameric receptors, but homomeric receptors usually comprise subunit α5 or α7 only, and other subunits can only form heteromeric receptors. At the same time, the heteromeric receptors, in contrast to the homomeric counterparts, possess two or more orthosteric ACh-binding sites at the junction between the primary (+) face of an α subunit and the complementary (−) side of a β subunit. In addition, receptors including α4 usually are more sensitive to nicotine [10–11], and the homomeric receptor containing α7 not only is very sensitive toα-bungarotoxin, but also has high calcium ions permeability [12–14]. Thus, detecting the sites specific to each subunit, is important to understand the properties of the subunits, as well as the receptors formed by them.

According to the neutral theory of evolution [15], some amino acid sites in a protein may change in evolution without affecting the general function of the molecule, but a few sites subject to strong evolutionary constraints are much conservative, and substitutions on these sites may cause modification in protein structure or function. It is known that in many protein families, the specific functional characteristics of the members are determined by a small subset of amino acids at particular positions. Identification of such functional specific sites is critical for studying these proteins, such as function prediction and drug design. In recent years, different algorithms have been developed for pinpointing protein sites responsible for functional specificity. These algorithms usually are based on relative entropy, evolutionary rates, and the physical and chemical properties of amino acids in the sequences [16]. In a previous study, we conducted a phylogenetic analysis on nine neuronal nAChR subunits and confirmed their homology [17]. In the current work, we employed the evolutionary trace (ET) method [18] and the two-state model [19] to detect and investigate the functionally crucial sites within nAChR subunits. The ET approach relies on identifying conserved amino acid patterns in sequence alignments to highlight functional epitopes and essential residues for ligand binding specificity.

By integrating mutational evolutionary analysis and structural similarities within protein families, the ET method offers an evolutionary perspective for evaluating the functional or structural significance of individual residues within the protein’s structure. On the other hand, the two-state model categorizes sites into four groups and distinguishes radical and conserved substitution sites based on the physical and chemical characteristics of amino acids. This approach can establish a framework for protein functional evolution by combining the biochemical properties of amino acids with sequence evolution models.

By analyzing the amino acid sequences of neuronal nAChR subunits expressed in human and other vertebrate brain, we detected potential functional specific sites in each subunit, which could provide useful clues to understand their functional differences.

## Materials and methods

### Identification of functional conserved sites

Based on results of sequence alignment and phylogenetic analysis of 105 protein sequences in 9 neuronal nAChR subunits of 12 vertebrates [9](Supplemental Table S1), the ET method was used to identify the functional conserved sites in the nine neuronal nAChR subunits. ET uses phylogenetic analysis to identify amino acid sites important to protein structure or function based on alignment of homologous amino acid sequences [10].The idea of this method is to systematically define family subgroups of similar sequence (and, by implication, of similar functional expression) with the aid of phylogenetic tree, and then identify those residue positions that are invariant within each subgroup. These sites are conserved in subgroups involution, implicating that they may be vital to some shared characteristics of the subgroups [12]. With the increase of branching differences, the correlation between branching sites and functional differences between proteins became stronger [13].

The steps of ET to identify functional conserved sites are as follows: (1) the phylogenetic tree of homologous proteins was constructed by multiple sequence alignment; (2) partition identity cutoffs (PICs) are used to define partitions of phylogenetic tree; (3) each partition can generate a consensus sequence, and if the site has not changed within the group, the type of its conserved residue is assigned, otherwise it is left blank. At low PIC, sequences of group vary widely. At high PIC, proteins in the same group are increasingly similar in sequence and function. Finally, the tracked functional sites were mapped to the tertiary structure of the protein, and the spatial positions of amino acid sites were combined for further analysis [10, 12].

We used TraveSuite platform [14] to construct a phylogenetic tree using Kitsch algorithm and divide it into 10 partitions. Then, TraceSeq and TraceScript algorithms were used to track the site in each partition [15]. Finally, we use JDet (http://csbg.cnb.csic.es/JDet/) to represent the identified sites [16].

### Identification of sites related to functional divergence

Differences in function between family members after gene replication are usually accompanied by changes in the rate of sites evolution [17–19]. According to Gu et al., there are four types of amino acid sites for two duplicate gene clusters, i.e., type 0 sites are generally conserved throughout the gene family; type Ι sites are conservative in one cluster and variable in another; type II sites are conservative in both clusters, but with large functional differences; type U sites are variable in both clusters.

Some specific conserved sites different from other subgroups were not directly related to functional differences. In the standard of site mutation experiment, if the physical and chemical properties of amino acids change greatly after the site mutation, it is likely to have a greater impact on the functional change [20]. Based on the physicochemical properties of amino acids, we screened the sites of type functional divergence, *θ*_I_, then calculated the coefficient of type functional divergence between subunits using two-state model and maximum likelihood method, and calculated the correlation values with functional divergence for different amino acid substitution sites [11, 17, 21]. The two-state model assumes that the site may experience two states early in gene duplication, *F*_0_ (type II unrelated) and *F*_1_ (type II related). The probability of a residue being under *F*_1_ is P(*F*_1_) = *θ*_I_ and that being under *F*_0_ is P(*F*_0_)=1-*θ*_I_. In the later stages, an amino acid residue is always under the state of *F*_0_, amino acid substitutions in this stage are mainly under purifying selection.

In this study, we used DIVERGE 3.0 to calculate *θ*_I_, and Z-test was used to identify whether the two gene subgroups had obvious type functional divergence. For the subgroup with obvious functional divergence, Bayesian algorithm is used to calculate the posterior probability and predict the sites of type functional divergence. Compared with ET, DIVERGE divides the amino acids into four groups according to their physical and chemical properties, i.e., charge positive (K, R, and H), charge negative (D and E), hydrophilic (S, T, N, Q, C, G, and P), and hydrophobic (A, I, L, M, F, W, V, and Y). An amino acid substitution is called radical if it changes from one group to another; otherwise it is called conserved, that is, within the group [11].

### Mapping of sites to ***α***7 tertiary structure

The Cryo-EM structures of the a7 nicotinic acetylcholine receptor was recently determined and the structure information was acquired from the Protein Data Bank (PDB; https://www.rcsb.org; ID: 7KOQ)[22]. The identified sites were projected onto the A chain of α7 using PyMol (https://pymol.org/) to observe the relationship between site function and spatial position.

## Results

### Conserved sites in the vertebrate neuronal nAChR subunit

The phylogenetic tree of neuronal nAChR subunits were divided into 10 partitions using the ET method (Supplementary Figure S1). In the consequent analysis, we tracked the three partitions of P01-P03 and analyze the identified sites.

A total of 55 completely conserved sites in the vertebrate neuronal nAChR subunits were traced by P01 partition (Supplementary Figure S2). Of these sites, 29 were distributed in ECD, 24 sites were distributed in TM1-TM3, and 2 sites (Val290 and MET305) were located in the α helix region of the first segment of ICD connecting TM3. Projecting these sites onto the tertiary structure of α7, three groups of densely distributed sites were identified (Figure 1A)[23]. The sites of the first group (Figure 1B) were mainly distributed on loops A-C, β2-β3, β3-β4 linker, which overlapped with the positive binding interface of ligand binding domain (LBD) in space. The second set of sites (Figure 1C) was located between the LBD and the ion channel, of which 7 sites were distributed in the Cys-loop (Cys128-Cys142). Ala96, located on the β4-β5 linker, and Lys125, located on β6, were sequentially distant but closely linked in tertiary structure. A salt bridge was formed between the residues of Asp138 and Arg206, thus connecting the conformational transfer between LBD and ion channel [24]. The third group of sites was distributed in the transmembrane helical region of TM1-TM3. Although the sites of TM1 and TM3 were far apart in sequence, they were closely connected in tertiary structure. Notably, for Phe230 on TM1, Phe253 on TM2, and Met279 on TM3, side-chain benzene rings or sulfur atoms were close to residues on other TM segments, resulting in stable chemical structures (Figure 1D). As can be seen from the distribution of these three groups of sites, the positive binding pocket of LBD, TM1-TM3 transmembrane helix, and the energetic coupling pathway between LBD and ion channel remained stable in the evolution of vertebrate neuronal nAChR [20].

**Figure 1.**
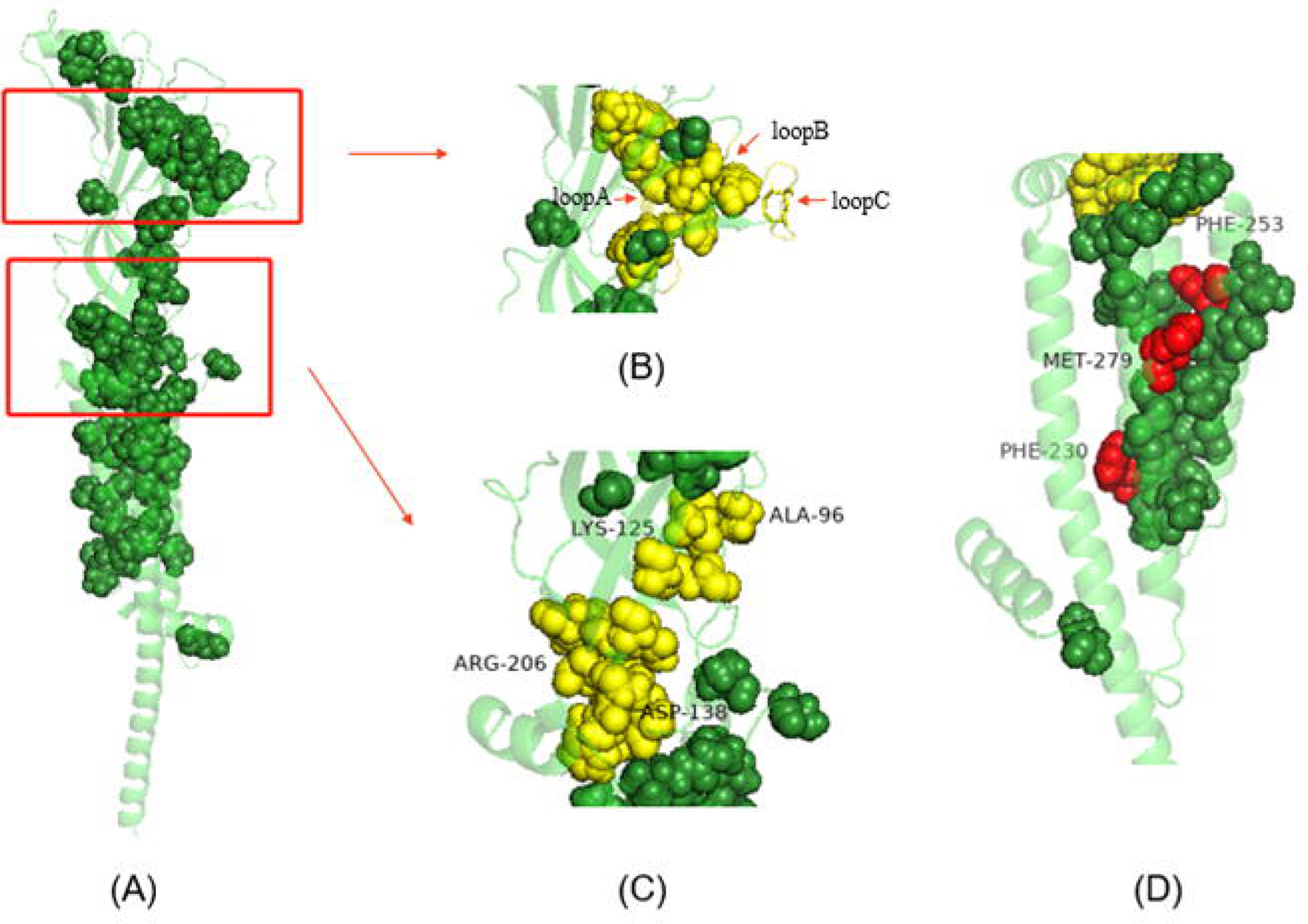
Spatial distribution of functional conserved sites of nine neuronal nAChR subunits in vertebrates. (A) 55 completely conserved sites tracked by P01 were mapped to α7 tertiary structure. (B) The first set of sites (Trp60, Trp86, Pro88, Asp89, Leu92, Phe146, Ser148, Trp149, Tyr151, Tyr195), loopsA-C, L92 at loopA, F146 at loopB, and disulfide bonds between Cys on loopC are highlighted in yellow. (C) The second set of sites (Ala96, Lys125, Ser126, Cys128, Phe135, Pro136, Phe137, Asp138, Gln140, Cys142, Glu173, Trp174, Arg206, Tyr211) located in LBD and TMD were highlighted.Interlinked Ala96 and Lys125,Asp138 and Arg206 are labeled.(D) By highlighting Phe230, Phe253 and Met279, it is speculated that these sites play an important role in the connection of the three TM regions.

### Conserved sites specific in ***α***7 subunit

P02 tracked 22 specific conserved sites in the α7 subunit, as well as in other subunits, but different from the α7 subunit (Supplementary Figure S2). These sites were distributed in ECD, TM1-TM3, C-terminal and the α helix region of the first segment of ICD connecting TM3 (Figure 2A). The β4-β5 linker connected to loopA at the sequence site and was spatially adjacent to the β6’ fold, which continued loopE in the sequence. In the case of nicotine molecule binding, loopA was related to loop E being an LBD pocket. Four sites of the α7 subunit were located in ECD, and Tyr118 was located at loop E, adjacent to the conserved Leu119, which was conserved non-polar tryptophan in other subunits. Ser95 was attached to Glu98 on the β4-β5 linker, and Glu98 was spatially closely linked to Lys125 on β6’ (Figure 2B). In other subunits, Glu98 was replaced by polar glycine, which attenuated the intersite forces (Figure 2C). The weak acid Thr208 and Tyr210 at the entrance of α7 subunit were replaced by non-polar proline and phenylalanine, which probably attenuated the negative charge environment of ion channel on the surface of cell membrane, and was related to the decrease of calcium ion permeability. The α7 subunit was highly conserved in the α helix region of the first segment of ICD connected to TM3, which was specific to the α7 subunit. We found that this structure was important for the expression and function of the α7 subunit [25]. P469 and N470 distributed at the C-terminus of the α7 subunit were space-aligned in other subunits. These two sites can also be regarded as specifically conserved in the α7 subunit and may be involved in the interaction between TM4 and the cell membrane.

**Figure 2.**
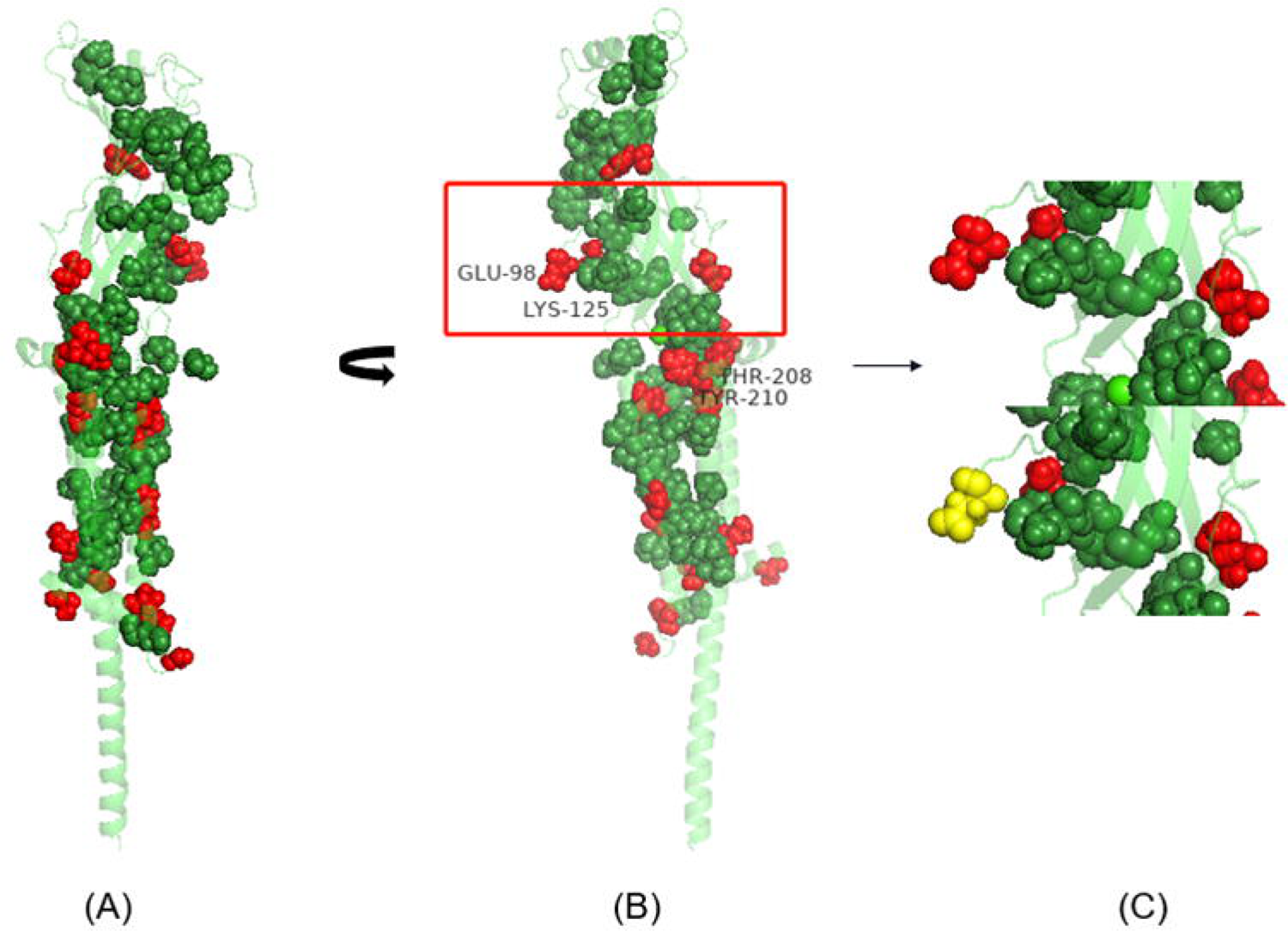
Spatial distribution of α7 subunit specific conserved sites identified by P02. (A) The 55 completely conserved sites identified by P01 were highlighted in green, and the 20 α7 subunit specific conserved sites identified by P02 are highlighted in red. (B) Rotate figure A by an Angle. (C) Glu98 conserved in the α7 subunit is replaced with glycine conserved in other subunits, and the substituted glycine is highlighted in yellow.

### Conserved sites specific to subunit subsets

P03 traced a total of 13 sites, which identified α2-α6 and β3 subunits as group 1, β2 and β4 subunits as group 2, and α7 subunits as group 3. These sites were conserved in each of the three groups. Asn107 and Ala226 in group 3 were replaced by basic lysine and polar threonine in group 1. Asn107, located in β5 and adjacent to loopE, was polar amino acid, asparagine, in both α7 and β2β4 subunits. After the second evolution, it was replaced by conserved alkaline lysine in the first group. In fact, when nicotine was bound to the ligand, the hydrophobic properties of the negative subunit loop E were more conducive to the binding of the ligand, which explained that nicotine can be bound to the nicotine molecule as a negative binding interface. However, the α4 subunit replaced the three hydrophobic subunits of loop E by one basic amino acid and two polar amino acids, thus rejecting ligand binding [26].

The conserved sites in the second group corresponded to Phe100, Asp131, Gly147, Ser223, and Ser249 in the α7 subunit. The non-polar amino acid Phe100 was located in the β4-β5 linker, adjacent to 1oopA (Leu92). Because Tyr93 of loop A was an important site for binding ligand molecules such as nicotine, the non-polar aromatic amino acid was replaced by the weak acid polar aromatic amino acid at Phe100. It was likely to affect the ligand binding of loopA or the structural changes after ligand binding. The polar amino acid Gly147 at loop B corresponds to positively charged arginine in the second group. It has been shown that in α4β2 receptors, if the β2 subunit acts as a positive binding interface, the arginine group at this site will be wedged to loop C and loop A at the β2 subunit, thus occupying the binding site of the nicotine ligand molecule.

Asp131 was located in the Cys-loop region and plays an important role in energy transfer from LBD to ion gated channels. Arg133, Met261, Ala263, Ser267, Gln273 and Ile281 in the α7 subunit were conserved in the three groups, but not in different groups. Arg133, a basic amino acid located in the Cys-loop region, was a conserved basic amino acid in the second and third group; at the same site in the first group, amino acid threonine was presented, which was presumed to play different regulatory role in the structural changes of energy transfer. Lys274 was the site with the greatest change in the physical and chemical properties of amino acids between the three groups, located at the entrance of TM3 and adjacent to the spatial location of Cys-loop. The site in this region was related to the conformational transmission of LBD to ion channel, as well as the charge accumulation near the cell membrane [27].

### Functional divergence sites between α7 subunit and other neural subunits

We calculated the functional divergence coefficients of type among nine nAChR pairwise groups. The functional divergence coefficients of type among α2α4, α3α6 and β2β4 were not statistically significant, but the functional divergence coefficients of other subunits were significant. Notably, all subunits had the highest degree of type divergence relative to the α7 subunit (Table 1). We calculated the posterior probability ratio of sites of functional divergence between the α7 subunit and other neuronal subunits. Based on the posterior probability ratio of sites, two classes of sites conserved within the α7 subunit and in one or more of the other subunits, were identified. Compared with α7 subunit amino acids, the first type of sites had radical substitution, and the second type of sites had conserved substitution. Altogether, 74 substitution sites were identified between α7 and other subunits, which were mainly distributed in β4-β5 linker, Cys-loop, β10-TM1 linker, TM2-TM3 loop, and the initiation region of ICD (Table 2). These sites were likely involved in functional differences relative to the α7 subunit.

**Table 1.** Type L functional divergence coefficients of 9 nAChR subunits in 12 vertebrates (*θ*_I_ ±SE)

| | $\alpha 7$ | $\alpha 2$ | $\alpha 3$ | $\alpha 4$ | $\alpha 5$ | $\alpha 6$ | $\beta 2$ | $\beta 3$ |
| --- | --- | --- | --- | --- | --- | --- | --- | --- |
| $\alpha 2$ | $0.43 \pm 0.04^*$ | | | | | | | |
| $\alpha 3$ | $0.46 \pm 0.04^*$ | $0.12 \pm 0.04^*$ | | | | | | |
| $\alpha 4$ | $0.45 \pm 0.04^*$ | $0.00 \pm 0.04$ | $0.11 \pm 0.04^*$ | | | | | |
| $\alpha 5$ | $0.52 \pm 0.04^*$ | $0.24 \pm 0.04^*$ | $0.29 \pm 0.04^*$ | $0.24 \pm 0.04^*$ | | | | |
| $\alpha 6$ | $0.45 \pm 0.04^*$ | $0.15 \pm 0.05^*$ | $0.06 \pm 0.05$ | $0.13 \pm 0.05^*$ | $0.25 \pm 0.05^*$ | | | |
| $\beta 2$ | $0.49 \pm 0.04^*$ | $0.26 \pm 0.04^*$ | $0.29 \pm 0.04^*$ | $0.27 \pm 0.04^*$ | $0.40 \pm 0.04^*$ | $0.32 \pm 0.05^*$ | | |
| $\beta 3$ | $0.49 \pm 0.04^*$ | $0.24 \pm 0.04^*$ | $0.29 \pm 0.04^*$ | $0.23 \pm 0.04^*$ | $0.08 \pm 0.05^\#$ | $0.26 \pm 0.05^*$ | $0.37 \pm 0.04^*$ | |
| $\beta 4$ | $0.42 \pm 0.04^*$ | $0.20 \pm 0.05^*$ | $0.24 \pm 0.05^*$ | $0.19 \pm 0.04^*$ | $0.33 \pm 0.05^*$ | $0.29 \pm 0.05^*$ | $0.02 \pm 0.04$ | $0.29 \pm 0.05^*$ |

$\theta_i$ is given by the maximum likelihood method, and SE represents the standard deviation of $\theta_i$ .
P value is calculated according to Z test of $(\theta_i / \text{SE})$ , where \* represents $p < 0.005$ , # represents $0.01 < p < 0.05$ , indicating the difference between the two subunits is statistically significant.

**Table 2.**
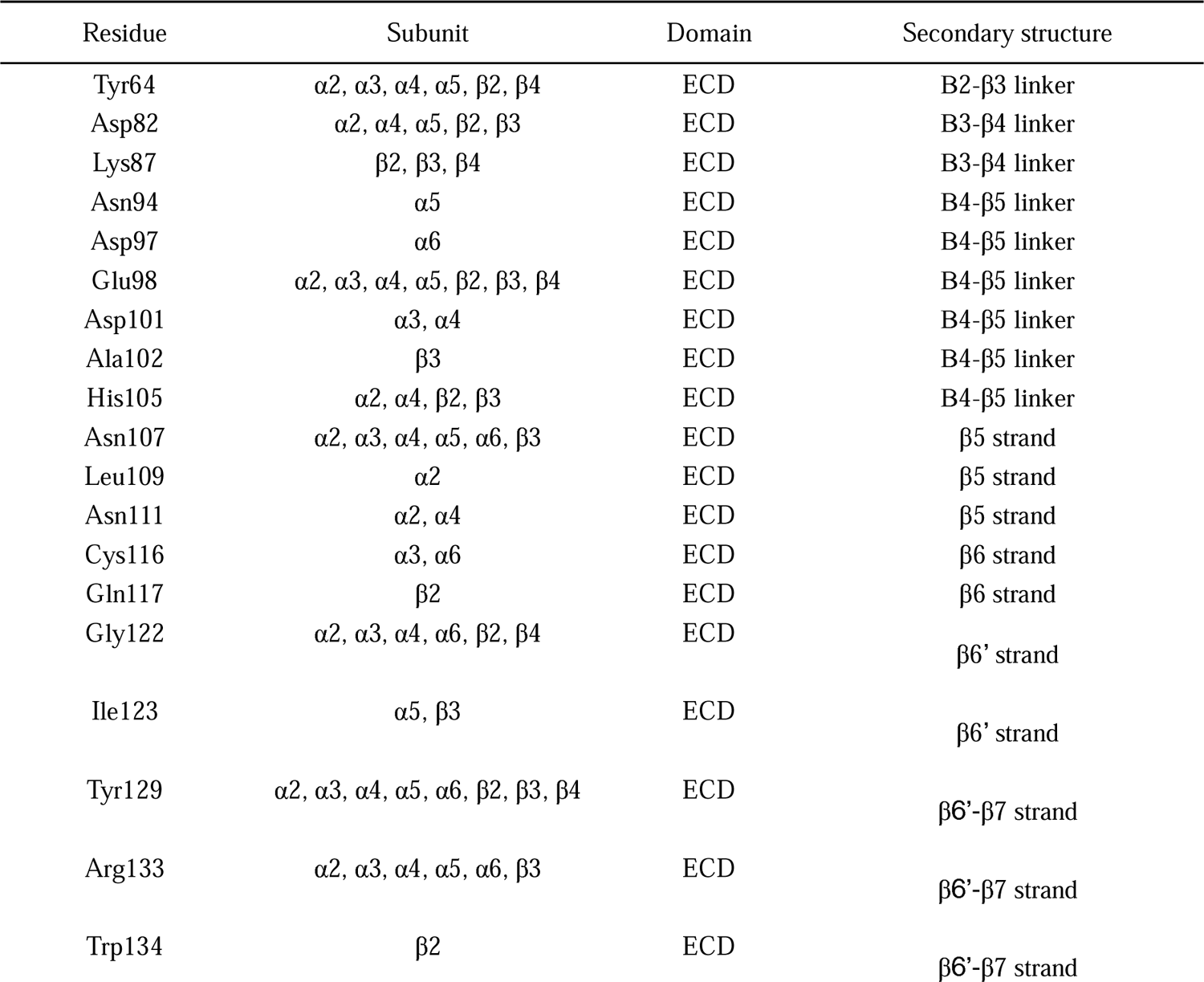
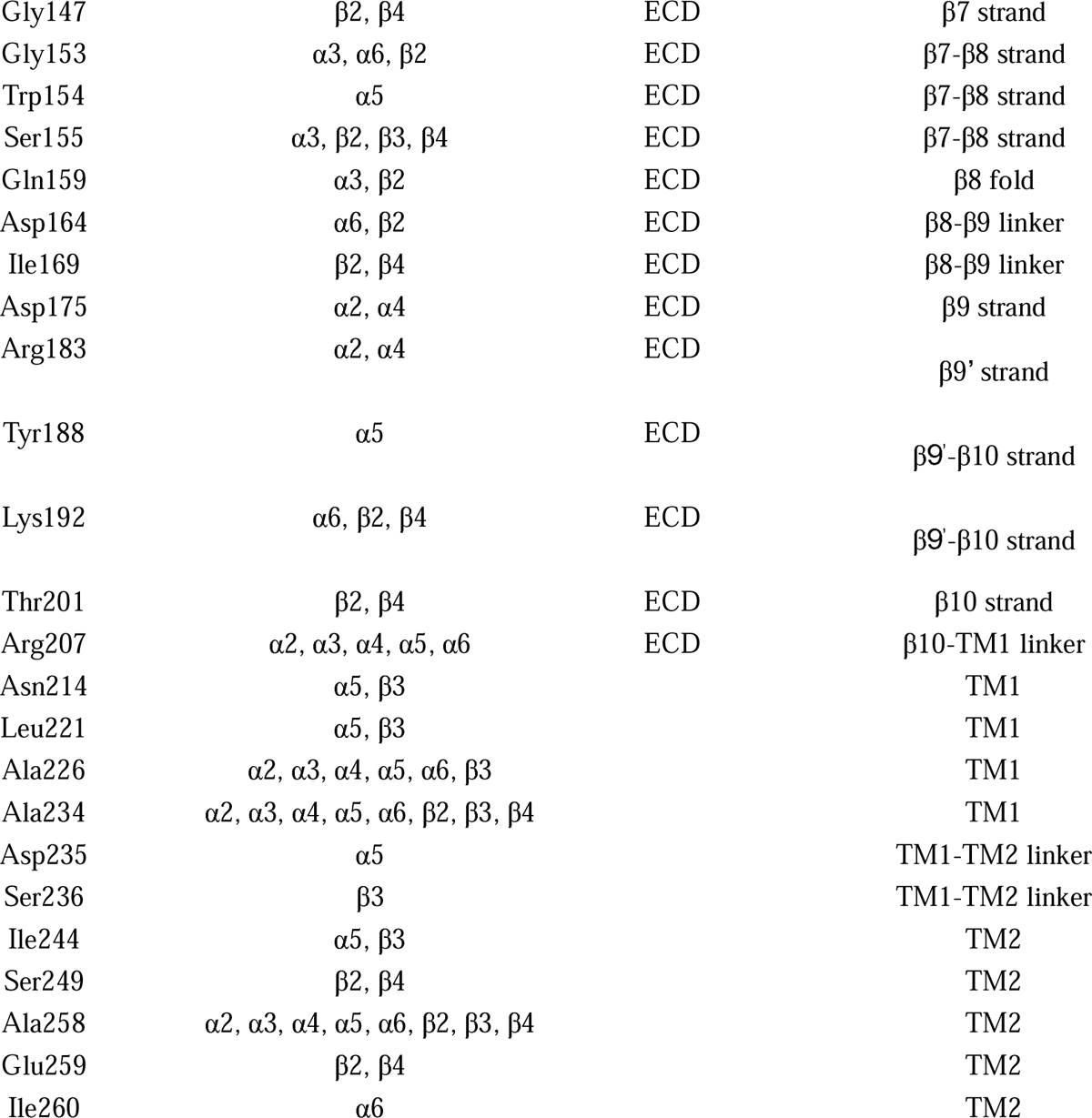
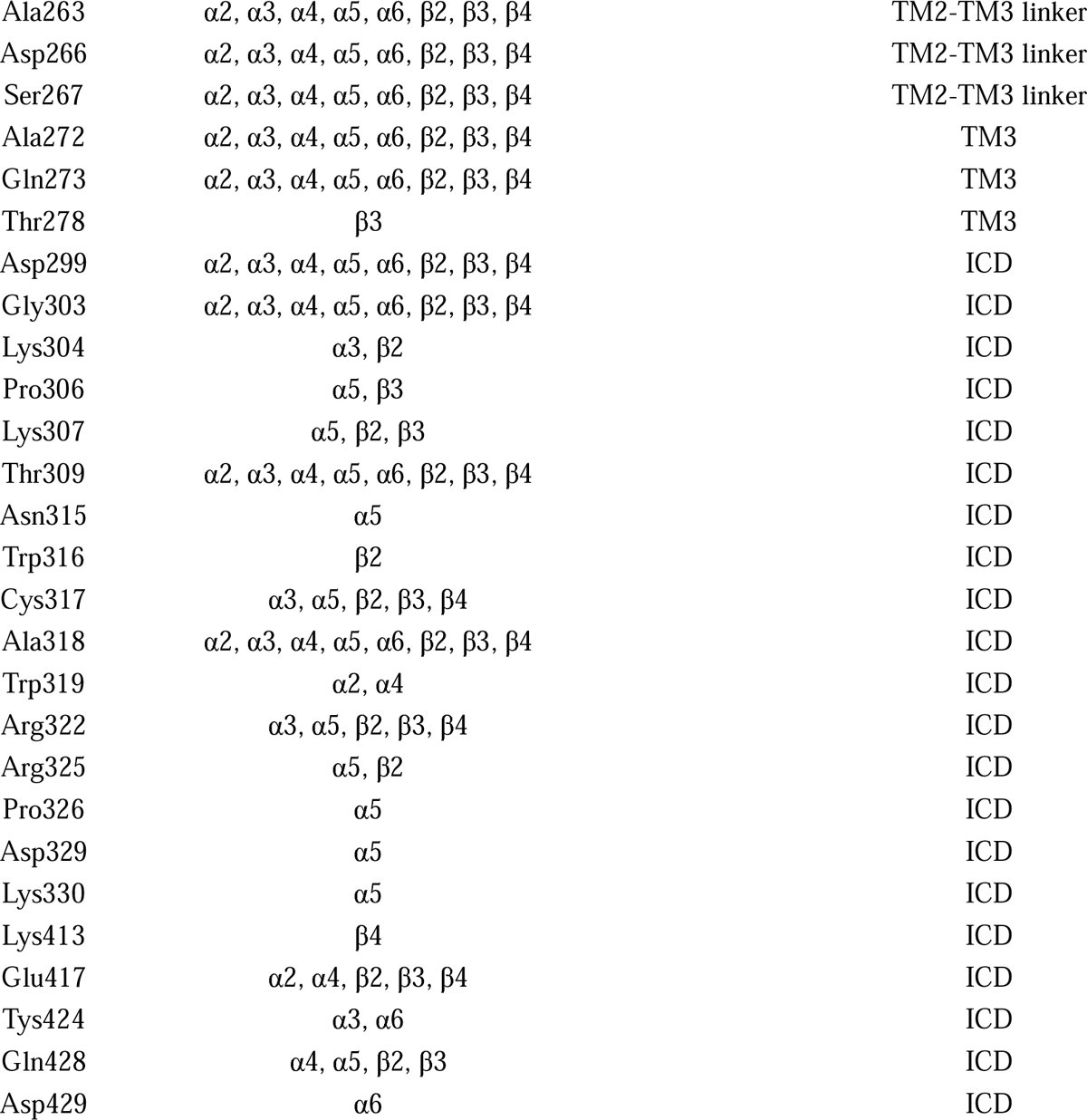
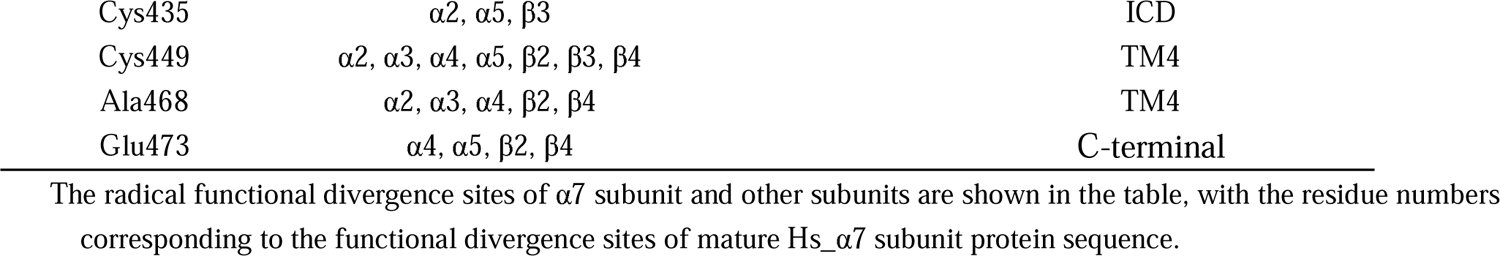
Functional divergence sites of intercomparison between each subunit and α7 subunit.

By mapping 74 class I sites to α7 tertiary structures and combining with 55 fully conserved sites tracked by ET, we can see that the fully conserved sites of subunits are mainly distributed in the interior of the receptor, while the new conserved functional sites that are easy to emerge in evolution are mainly distributed on the surface of the receptor (Figure 3A). There are eight sites (Tyr129, Ala258, Ala263, Asp266, Ser267, Gln273, Asp299, Gly303) that are conserved within each subunit but have altered amino acid properties relative to the α7 subunit (Figure 3B). Cys-loop is adjacent to the TM2-TM3 loop in spatial position, and plays an important role in the energy transfer from LBD to ion channel. Tyr129 is located in the Cys-loop, while Asp266 and Ser267 are located in the TM2-TM3 loop. Asp266 is replaced by lysine in the α5β3 subunit and leucine in the other six subunits. Asp266 was used to distinguish the three groups of subunits. The functional difference lies in that the acidic α7 subunit can form homomers at this site, while the non-polar subunit can only form heteromers in pairs. The basic α5 and β3 subunits at this site do not participate in ligand binding. Arg133 on the Cys-loop is a basic lysine in the β2β4 subunit and a polar threonine in the α2-α6 and β3 subunits, which may also be related to the fact that the three subunits provide negative binding interfaces (Figure 3C).

**Figure 3.**
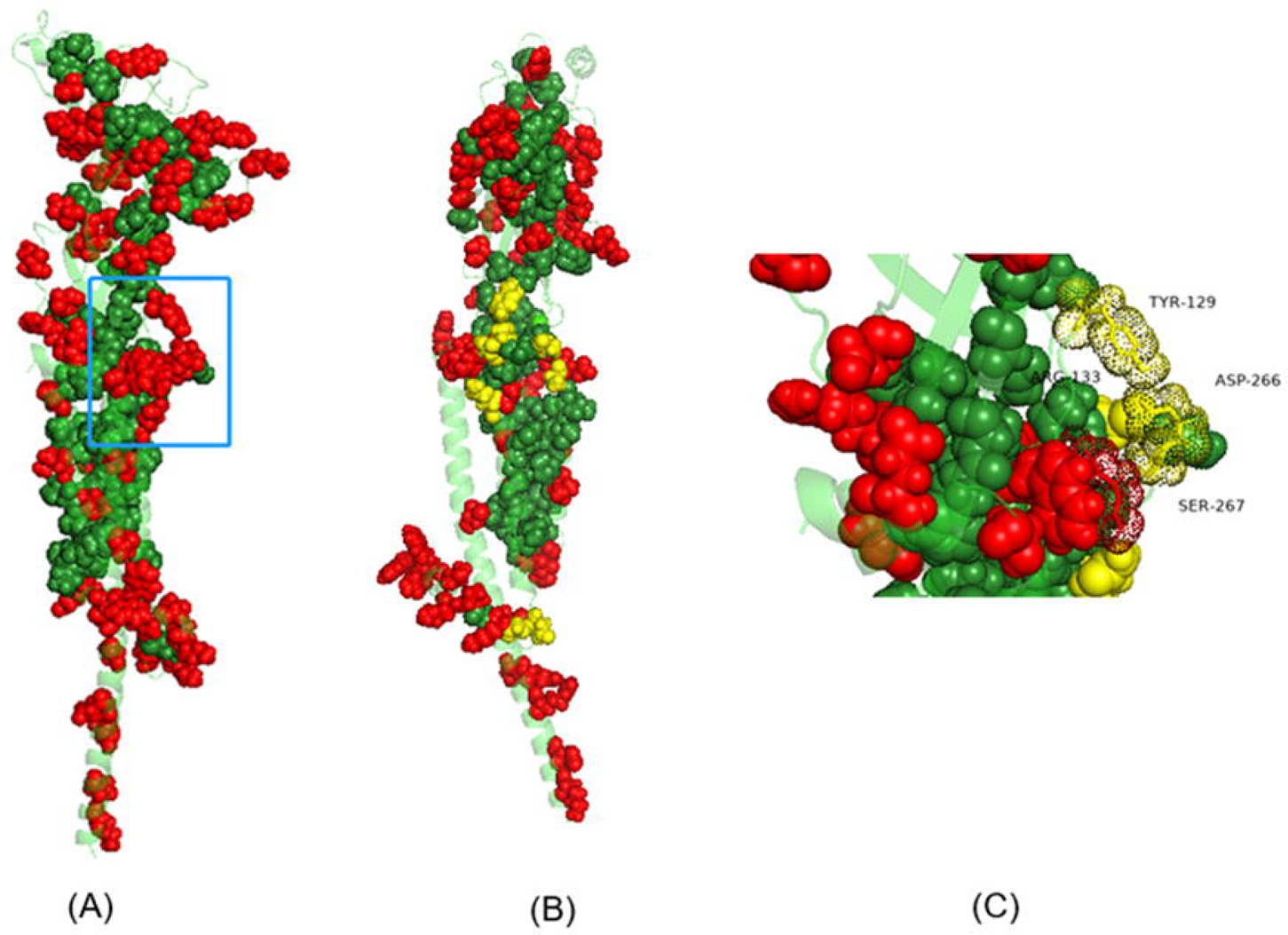
Spatial distribution of conserved sites of total subunit and specific functional divergence sites of α7 subunit. (A) The completely conserved sites measured by ET were highlighted in green, and the specific functional divergence sites of the α7 subunit were highlighted in red. (B) Figure (A) was rotated to indicate yellow sites that were conserved in each subunit but changed in physicochemical properties relative to the α7 subunit., C-The area in the blue box in Figure (A), the detail between the “Cys-loop” and the TM2-TM3 loop.

### Functional divergence sites in subunit subsets

By calculating the posterior probability ratio of functional divergence sites of α7 subunit to other 8 subunits, 5 qualitative variant substitutions and 15 conservative substitutions were obtained, which were similar to those traced by P02 partition of ET (Supplementary Figure S3). Compared with ET, this method classifies the sites tracked by P02 and quantitatively calculates the posterior probability ratio. The sites where the properties of the first group of amino acids changed are more likely to be related to differences in the basic biological functions of the two subunits.

According to the posterior probability ratio, a total of 29 radical substitution sites and 43 conserved substitution sites were obtained between α7 and β2β4 subunits. These sites include all sites tracked by P02 partition of ET except c-terminal and some sites tracked by P03 partition. The functional divergence sites between α7 subunit and β2β4 subunit were mainly distributed in “Cys-loop”, TM2-TM3 loop, loopC and LBD binding interface near the connected linker region. Therefore, it can be speculated that the two groups of subunits have great functional differences in the selective binding ability of ligand and the response coefficient of ion channel after ligand binding.

There were 11 functional divergence sites between α7 subunit and α2-α6 and β3 subunit groups, all of which were identified in the P03 partition of ET, and no functional divergence sites of conserved substitution were found. Among the functional disambiguation sites between β2β4 and α2-α6 and β3 subunits, only I291 in TM3 was not identified in RT-P03. Although the posterior probability ratio of Glu259 was low, multi-sequence alignment revealed that the basic amino acids in β2β4 subunit were conserved, and the acidic amino acids in α3-α7 and β3 subunit were conserved, and the replacement of acidic amino acids only occurred in α2 subunit. This acid-base difference was likely related to the characteristics of receptor channels.

Finally, we divided α2α4, α3α6 and α5β3 subunits into three groups. The posterior probability values of α2α4 and α5β3, α3α6 and α5β3 showed significant functional divergence. The functional divergence sites were distributed in TM3 linker, ICD, β7β8 and β9’. It can be seen that the large functional differences among the three subunits are related to the characteristics of ion channels.

## Discussion

As the major receptors in cholinergic system, nAChRs play important roles in a series of physiological processes. They are also involved in neurological disorders like nicotine addiction, Alzheimer’s disease and Parkinson’s disease [28–31]. Thus, these complexes are promising targets of therapeutic agents [32–34]. The diversity in function and types of nAChRs relies on variety of properties of the subunits and their combinations. Since the neuronal subunits in vertebrates were descended from common ancestors, comparing these subunits in the context of evolution will provide valuable clues to understand their specificity.

By using ET, we identified 55 completely conserved sites in nine vertebrate nAChR subunits. These sites are mainly distributed in the positive binding surface of ligand binding domain, the input interface region of “LBD-TMD energy pathway” (β8-β9 connection loop and Cys-loop), the outer membrane inlet and outlet end of transmembrane helices, and the interconnecting sites between three transmembrane helices. These regions are important for the basic structure or function of the protein family. Previous experiments have shown that Loop A-C and Loop D-F provide positive and negative binding interface for binding domain, and loopsD-F has higher sequence variability than Loops A-C [35–37]. The distribution region of one of the loci identified by P01 partition of ET overlapped with the positive binding interface, which verified that the positive binding interface played a stronger role in the binding of nAChR to the ligand. Compared with the positive binding surface, the negative binding surface is more suitable for the regulation of ligand binding parameters. The energetic coupling pathway between LBD and ion channel gated region confirmed by Lee WY [38] and the 7 highly conserved sites in the superfamily measured by Ortells [39] are consistent with the results obtained by P01, thus confirming the important role of energetic coupling pathway in nicotinic receptor channel protein [40]. In addition, β4-β5 is adjacent to β4 and β6’, β4 is a component of loop A, β6’ is a continuation of loop E, β4-β5 loop is approximately located in the middle of positive and negative binding interfaces in space, which may affect the conformation of the two binding interfaces.

Multiple sites on the TM2-TM3 loop, the outer end of TM2 membrane and the outer beginning of TM3 membrane have been proved to be important components of the energetic coupling pathway, participating in the conformational transfer of LBD to the gated region of the receptor protein ion channel. In addition, this region is related to the concentration of ions that the receptor accumulates around the cell membrane, thus affecting the conductivity and calcium permeability of the channel. For example, the site near the entrance of TM3 of α7 subunit is polar, but less polar than other subunits, similar to the intermediate state of β2β4 subunit and other subunits. It is inferred that the difference in this region is related to the better permeability of calcium ions, lower binding efficiency with nicotine molecules and faster desensitization of α7 subunit compared with other subunits. Through the spatial distribution of sites, we found that all the conserved sites in the α7 subunit were concentrated in the positive binding interface of LBD, the transmembrane helical region, the LBD-TMD energetic coupling pathway and the α helical segment in ICD cells that connected TM3. Finally, the total subunit conserved sites are mainly distributed in the inner region of protein spatial structure, while the functional divergence sites tend to be distributed in the surface region of protein surface. It is speculated that the subunits constantly generate new functions to adapt to changes in the external environment during the evolutionary process.

In summary, by using the statistical models such as evolutionary trace and the two-state model, we identified the conserved sites in nAChR subunits, and then analyzed the spatial distribution and their potential influence on the function of the subunits. The results showed that the conserved sites were concentrated in important regions related to ligand binding and ion channel energy transfer, and were closely related to the structure and function of the receptor. It is of great significance to analyze the sites of functional divergence among each subunit for understanding the functional differences among subunits, which will provide useful information for exploring the action mechanism of nAChR and targeted treatment of diseases.

## Acknowledgements

This study was supported in part by grants from the Joint Institute of Tobacco and Health (No. 2021539200340049). The funding bodies played no role in the design of the study and collection, analysis and interpretation of data, or in writing of the manuscript.

---

**Supplementary Figure S1.** Phylogenetic trees and evolutionary divisions of the nine neuronal nAChR subunits from 12 vertebrates. As PIC increases, from P01 to P10, partitions comprise more groups, each with fewer sequences.

**Supplementary Figure S2.** Functional conserved sites identified by ET. The site number corresponds to the amino acid site number of the α7 subunit protein sequence. (A) There were 55 completely conserved sites in P01 partition. (B) Specific conserved sites within the α7 subunit traced by P02. (3) Specific conserved sites traced by P03. The α4 subunit sites represent highly conserved in the entire first group, the β2 sites represent highly conserved in the entire second group, and the third group is α7.

**Supplementary Figure S3.** Functional divergence sites within subunit subsets identified by DIVERGE. The overlapping sites identified by ET were highlighted in red. The site number corresponds to the amino acid site number of the α7 subunit protein sequence. (A)Functional divergence sites between α7 subunit and other 8 subunits. (B) Functional divergence sites between α7 subunit and β2β4 subunits. (C) Functional divergence sites between α7 subunits and α2-α6, β3 subunits. (D) Functional divergence sites between β2β4 subunits and α2-α6, β3 subunits.

