## Supplementary figures and images for "Identification of the potential function-specific sites in subunits of vertebrate neuronal nicotinic acetylcholine receptors"

### Supplemental Figure S1

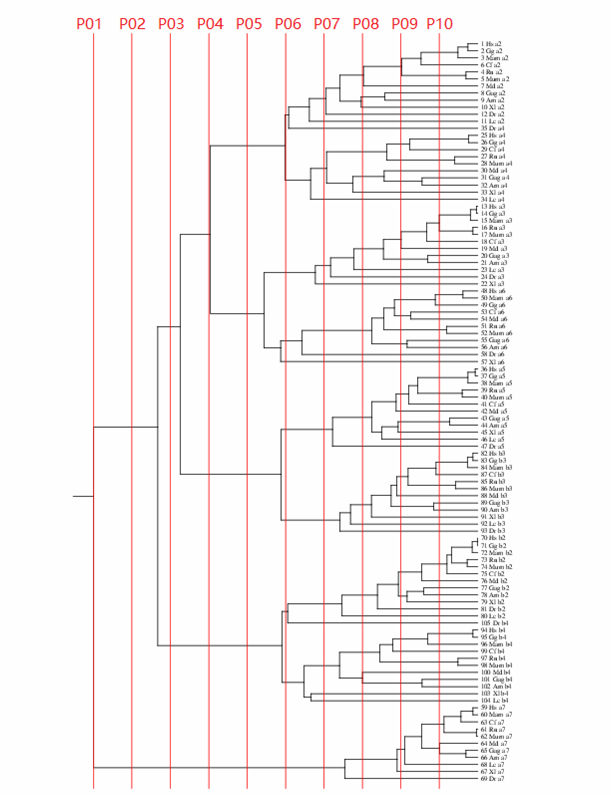

### Supplemental Figure S2

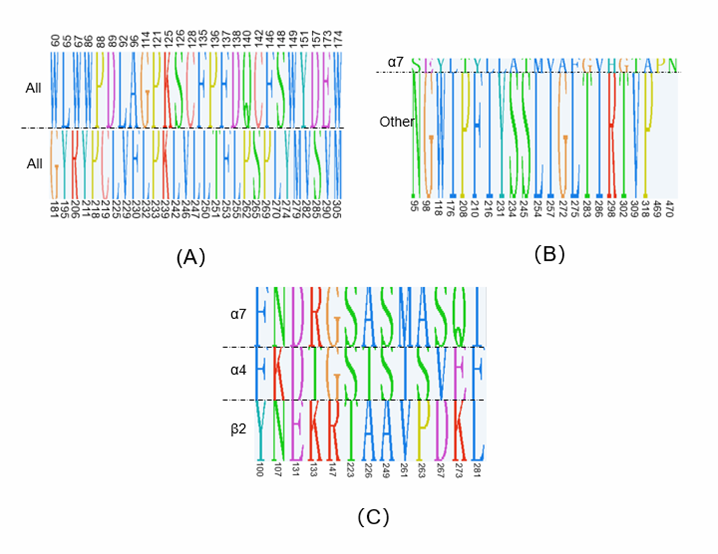

### Supplemental Figure S3

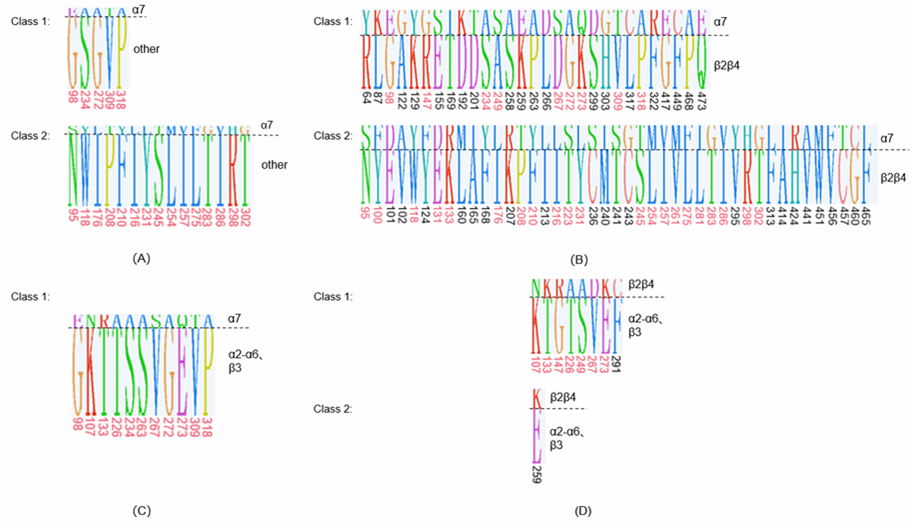
