## Supplemental Tables for "Identification of the potential function-specific sites in subunits of vertebrate neuronal nicotinic acetylcholine receptors"

**Supplemental Table 1**. Protein sequences of nine Neuronal nAChR subunits from 12 representative vertebrate species

| Gene Symbols | Species | Protein Accession Numbers |
| --- | --- | --- |
| CHRNA2 | H. sapiens | NP_000733.2 |
|  | G. gorilla | XP_004046867.1 |
|  | M. mulatta | EHH28371.1 |
|  | R. norvegicus | NP_596911.1 |
|  | M. musculus | NP_659052.1 |
|  | C. familiaris | NP_434692.2 |
|  | M. domestica | NP_660111.2 |
|  | G. gallus | NP_990146.1 |
|  | A. mississippiensis | XP_006274722.2 |
|  | X. laevis | XP_018119614.1 |
|  | L. chalumnae | XP_006010265.1 |
|  | D. rerio | NP_001035417.1 |
| CHRNA3 | H. sapiens | NP_000734.2 |
|  | G. gorilla | XP_004056656.1 |
|  | M. mulatta | XP_014998318.1 |
|  | R. norvegicus | NP_434692.2 |
|  | M. musculus | NP_660111.2 |
|  | C. familiaris | XP_013974315.1 |
|  | M. domestica | XP_007478065.1 |
|  | G. gallus | NP_989747.1 |
|  | A. mississippiensis | XP_006261416.1 |
|  | X. laevis | XP_018106913.1 |
|  | L. chalumnae | XP_006001903.1 |
|  | D. rerio | XP_001921314.1 |
| CHRNA4 | H. sapiens | NP_000735.1 |
|  | G. gorilla | XP_004062567.1 |
|  | R. norvegicus | NP_077330.1 |
|  | M. musculus | NP_056545.3 |
|  | C. familiaris | XP_013962793.1 |
|  | M. domestica | XP_001376203.1 |
|  | G. gallus | NP_990145.1 |
|  | A. mississippiensis | XP_019339959.1 |
|  | X. laevis | XP_018092533.1 |
|  | L. chalumnae | XP_005993532.1 |
|  | D. rerio | NP_001041528.1 |
| CHRNA5 | H. sapiens | NP_000736.2 |
|  | G. gorilla | XP_004056659.1 |
|  | M. mulatta | XP_014998319.1 |
|  | R. norvegicus | NP_058774.2 |
|  | M. musculus | NP_789814.2 |
|  | C. familiaris | XP_005628378.1 |
|  | M. domestica | XP_007478268.1 |
|  | G. gallus | NP_989746.1 |
|  | A. mississippiensis | XP_006261415.1 |
|  | X. laevis | NP_001086549.1 |
|  | L. chalumnae | XP_006001908.1 |
|  | D. rerio | NP_001017885.1 |
| CHRNA6 | H. sapiens | NP_004189.1 |
|  | G. gorilla | XP_004047018.1 |
|  | M. mulatta | XP_001099152.1 |
|  | R. norvegicus | NP_476532.1 |
|  | M. musculus | NP_067344.2 |
|  | C. familiaris | XP_539951.2 |
|  | M. domestica | XP_001381937.1 |
|  | G. gallus | NP_990695.1 |
|  | A. mississippiensis | XP_006277029.1 |
|  | X. laevis | XP_018108423.1 |
|  | D. rerio | NP_001036149.1 |
| CHRNA7 | H. sapiens | NP_000737.1 |
|  | M. mulatta | NP_001028055.1 |
|  | R. norvegicus | NP_036964.3 |
|  | M. musculus | NP_031416.3 |
|  | C. familiaris | XP_545813.4 |
|  | M. domestica | XP_001377515.2 |
|  | G. gallus | NP_989512.2 |
|  | A. mississippiensis | XP_014455380.1 |
|  | X. laevis | XP_018108505.1 |
|  | L. chalumnae | XP_006011750.1 |
|  | D. rerio | NP_957513.1 |
| CHRNB2 | H. sapiens | NP_000739.1 |
|  | G. gorilla | XP_004026880.1 |
|  | M. mulatta | XP_014965404.1 |
|  | R. norvegicus | NP_062170.1 |
|  | M. musculus | NP_033732.2 |
|  | C. familiaris | XP_547565.2 |
|  | M. domestica | XP_001373153.1 |
|  | G. gallus | NP_990144.1 |
|  | A. mississippiensis | KYO27011.1 |
|  | X. laevis | NP_001088299.1 |
|  | L. chalumnae | XP_005996559.1 |
|  | D. rerio | XP_005169811.1 |
| CHRNB3 | H. sapiens | NP_000740.1 |
|  | G. gorilla | XP_004047017.1 |
|  | M. mulatta | XP_015000759.1 |
|  | R. norvegicus | NP_598281.1 |
|  | M. musculus | NP_775304.1 |
|  | C. familiaris | XP_539952.3 |
|  | M. domestica | XP_007476474.1 |
|  | G. gallus | NP_990143.2 |
|  | A. mississippiensis | XP_006277025.2 |
|  | X. laevis | XP_018098965.1 |
|  | L. chalumnae | XP_006007648.1 |
|  | D. rerio | NP_957514.1 |
| CHRNB4 | H. sapiens | NP_000741.1 |
|  | G. gorilla | XP_004056660.1 |
|  | M. mulatta | XP_014998309.1 |
|  | R. norvegicus | NP_434693.1 |
|  | M. musculus | NP_683746.1 |
|  | C. familiaris | XP_852548.2 |
|  | M. domestica | XP_007478059.1 |
|  | G. gallus | NP_990150.1 |
|  | A. mississippiensis | XP_014461433.1 |
|  | X. laevis | XP_018108341.1 |
|  | L. chalumnae | XP_006001904.1 |
|  | D. rerio | XP_696993.2 |
